# Inhibition of select actinobacteria by the organophosphate pesticide chlorpyrifos

**DOI:** 10.1101/2021.03.18.436105

**Authors:** Nathan D. McDonald, Courtney E. Love, Rushyannah Killens-Cade, Jason Werth, Matthew Gebert, Carolyn F. Weber, Christopher Nealon, Charles Sweet, Noah Fierer, Henry S. Gibbons

## Abstract

Organophosphorus compounds have an extensive history as both agricultural pesticides as well as chemical nerve agents. Decades of research have demonstrated numerous links between these chemicals and their direct and indirect effects on humans and other organisms. The inhibitory effects of organophosphate pesticides (OPPs) on metazoan physiology, are well-characterized; however, the effects of organophosphorus compounds on soil microbes - essential contributors to key agricultural processes - are poorly understood. Chlorpyrifos (CPF) is an OPP that is used globally for crop protection. Studies of CPF application to soils have shown transient effects on soil microbial communities with conflicting data. Here, we directly test the effect of CPF on a panel of 196 actinobacteria strains, examining the effects of CPF on their growth and *in vitro* phenotypes on solid media. Strains were grown and replica-plated onto media containing CPF or a vehicle control and grown at 28°C. CPF dramatically inhibited the growth of most strains and/or altered colony morphologies, with 13 strains completely inhibited by CPF. In disk diffusion assays with CPF, its degradation product 3,5,6-trichloropyridinol (TCP), malathion, parathion, monocrotophos and mevinphos, only CPF exhibited direct antimicrobial activity suggesting that the observed effects were due to CPF itself.

**IMPORTANCE:** Chlorpyrifos is a globally used pesticide with documented neurological effects on non-target organisms in the environment. Finding that chlorpyrifos can inhibit the growth of some soil microbes *in vitro* may have implications for the composition, stability, and health of the soil microbiome. Due to the importance of soil microbes to numerous biogeochemical processes in agricultural systems, additional investigations into the non-target effects of CPF on soil microbes are clearly needed.

## INTRODUCTION

The global pesticide era began in the mid-20^th^ century, ushering in marked improvements in the agricultural industry and beyond (1). The production of crops benefited from dramatically increased yields through the second half of the 20^th^ century (2). While many factors contributed to the surge in crop production, there is no doubt that the expanded use of herbicides and pesticides played an integral role. Beyond increased yields and reduced crop losses, pesticide use has also led to better control of vector-borne diseases (1). Despite these benefits, the use of pesticides does not come without associated risk. A growing body of evidence has demonstrated various off-target effects of many pesticides on humans, other animals, and the environment (3-5).

The organophosphate pesticides (OPPs) are a class of insecticides broadly used in commercial agriculture worldwide. The OPPs function by forming a covalent OPP-enzyme adduct that leads to irreversible inhibition of the enzyme acetylcholinesterase (AChE), which terminates synaptic transmission by degrading the neurotransmitter acetylcholine (6). When OPPs bind to AChE, the enzyme hydrolyzes the OPP, leaving a phosphoryl group in the active site, irreversibly inhibiting AChE. Organophosphate pesticides are a major cause of intentional poisoning worldwide with early exposure symptoms including fatigue, salivation, and convulsions, ultimately leading to death by asphyxiation in acute cases (7). The toxic effects of OPPs on humans and other animals has been well documented, and the Environmental Protection Agency banned the residential use of many OPPs in 2001 (8-14). Despite this ban, many of these pesticides (**Fig. 1**) are still widely used in agriculture (15).

**Figure 1.**
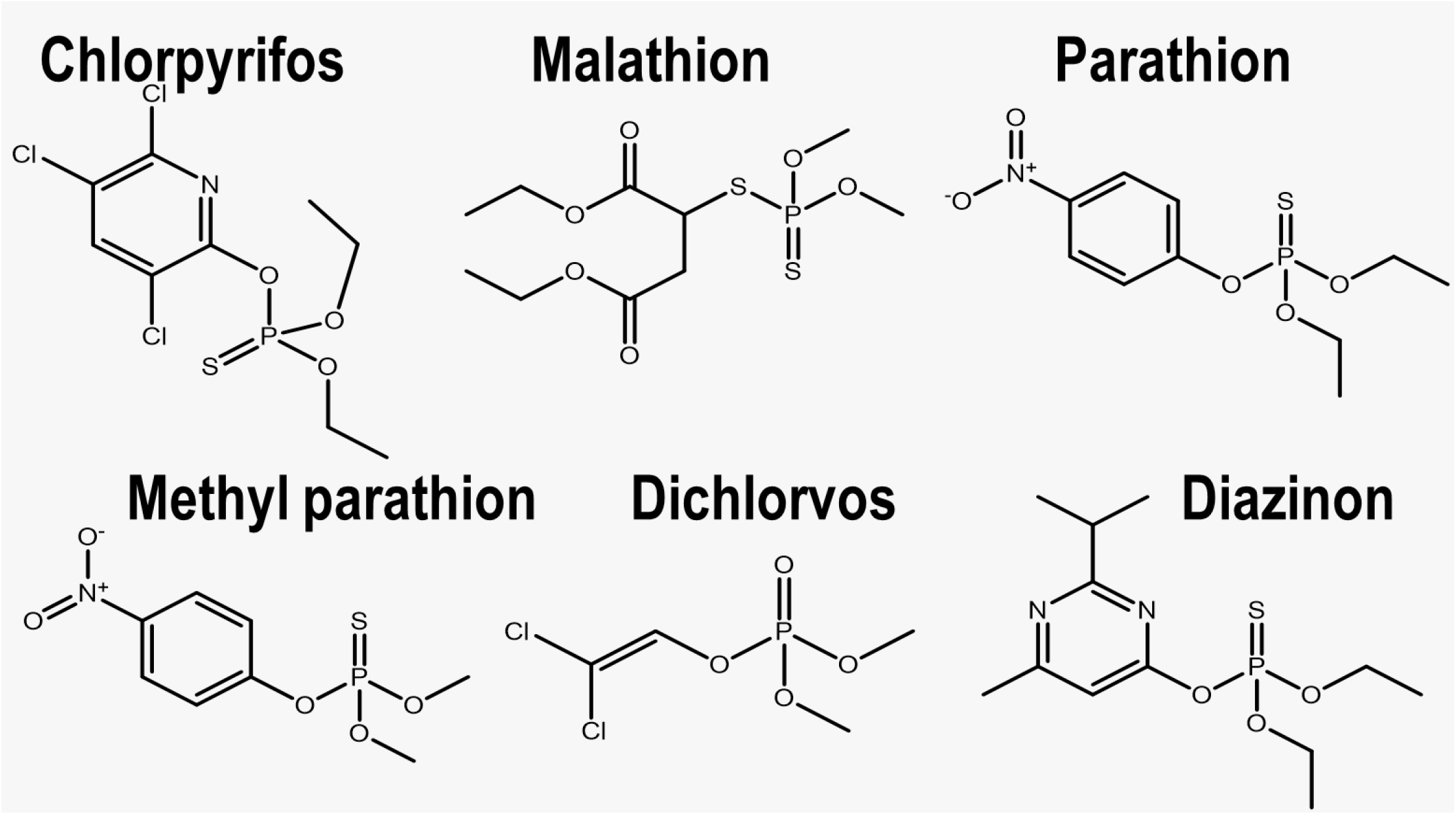
Representative organophosphate pesticides.

One of the most widely used and studied OPPs is chlorpyrifos (CPF) (**Fig.1**). Since its first introduction to the market in 1965, CPF has become the insecticide of choice for more than 50 types of crops in over 100 countries worldwide and is applied to 8.5 million crop acres per year (16, 17). Off-target exposure to CPF is associated with toxicity towards aquatic organisms (18-22), pollinators (23-26) and birds (18, 27-30). Additionally, several studies have shown a link between CPF exposure and adverse side effects on humans and other mammals (12, 31-38).

In contrast to the well-documented effects on human and animal health, the effects of OPPs on microbial populations following exposure to OPPs has received little study. Soil bacteria and fungi play an essential role in many agricultural processes, serving as key controls on soil nutrient cycling and other processes (39-42). It is well known that soil microbiomes contribute to overall soil fertility and crop yield (43-46). While there is limited and conflicting data on the effects of OPPs on microbes, several studies have investigated the possibility of utilizing bacteria as tools for CPF bioremediation (47-57). In fact, enzymes that degrade CPF, most notably organophosphate hydrolases, are present in various bacterial species (58-62). This bacterial degradation is believed to be one of the primary mechanisms by which CPF is removed from the environment (reviewed in (48)). While it is clear that there is interplay between soil microbes and CPF, the specific effects of this OPP and OPPs as a whole on soil microbes have not been well-studied. A few previous studies have demonstrated that exposure to CPF results in shifts in bacterial populations (48, 56, 63, 64); however, whether these shifts are due to inhibition of select microbes, enhanced growth, or a combination of both is less clear. It is also unclear if there are taxon-specific differences in bacterial responses to CPF.

Actinobacteria are one of the more abundant groups of bacteria found in soils across the globe (65). Actinobacteria are Gram-positive, high G+C content bacteria, many of which display characteristics of fungi such as production of differentiated multicellular structures (i.e., substrate mycelia, and aerial hyphae). While actinobacteria are predominantly found in soils, they can also be abundant in air and aquatic environments (66). Actinobacteria play significant roles in a variety of economic and environmental processes of importance. Actinomycetes affect crop production by fixing nitrogen, recycling organic matter, promoting plant growth, and in some cases, through plant pathogenesis (65, 66). (65, 66). Additionally, actinomycetes produce metabolites including antimicrobials, antifungals, and enzymes involved in bioremediation of numerous substances including OPPs (47, 57).

The purpose of this study was to characterize the effects of CPF on soil actinobacteria. Utilizing a panel of recently isolated actinobacteria, we demonstrate that the growth of many of the strains is delayed or completely inhibited following exposure to CPF. In addition, we noted several altered phenotypes including colony morphology and pigmentation. Using a modified Kirby-Bauer disk-diffusion assay, we demonstrate that inhibiton effects are specific to CPF, as its main degradation product 3,5,6 trichloropyridinol (TCP) and several other OPPs had no effect on bacterial growth. We next tested a panel of defined species across the actinobacteria obtained from the Agricultural Research Service (ARS) Northern Regional Research Laboratory (NRRL) culture collection for susceptibility to CPF. Across this panel we found that grwoth inhibition is specific to CPF, but not all species are equally affected and strains within the same species can exhibit resistant and sensitive phenotypes. This study provides evidence that CPF is able to inhibit the growth of specific actinobacterial taxa commonly found in soil.

## Results

### Chlorpyrifos exposure results in phenotypic changes in Actinomycetes

To determine whether CPF affects the growth of bacteria representative of taxa commonly found in soil ecosystems, we began with an array screen of 196 diverse strains within the family Actinomycetales grown in the presence or absence of CPF. The actinomycete panel represents strains/species which were isolated via air sampling across three geographical regions in the United States, including two sites in Maryland, one near Boulder Colorado., and one near Pocatello, Idaho (67). A final concentration of 10μM was selected because it would represent the effect of solubilizing the recommended field application of the pesticide in 2.54 cm of rainfall. Following the incubation, 144 of the 196 strains grew on the control plate which will serve as the reference (**Fig. 2**). Comparing the reference plates to the CPF plates we observed phenotypic changes in the presence of the OPP. Most notably, the majority of these observed phenotypes were related to growth inhibition (**Fig. 2**). Specifically, 44 strains (30.5%) exhibited delayed growth compared to the DMSO control. Delayed growth was scored by microbial growth on the CPF plates which was visibly reduced compared to the same strain on the reference plate. The library screen revealed that 33/144 (22.9%) strains were completely inhibited with no visible growth in the presence of CPF after the 3 day incubation compared to same strain on the control plate where growth was observed. Together these data show that, of the 144 strains that grew on the control media, a total of 77 (53%) were either partially or completely inhibited in growth by 10 μM CPF (**Fig. 2**) In addition to the effects on growth, CPF affected colony shape and coloration, which were likely due to delayed formation of aerial hyphae and/or sporulation (9/144; 6%) (**Fig. 2**). Taken together, these results indicate that, for individual species or strains, CPF at agriculturally relevant concentrations can have profound effects, with growth inhibition frequently observed. Of those CPF-responsive strains for which 16S rRNA gene sequences were obtained, all were species of *Streptomyces*. This finding is most likely due to the composition of the panel, which was originally intended to detect cryptic antimicrobial metabolites and as such contains a high proportion of Streptomyces spp., with minor representation (∼10%) from other actinomycete clades (Micromonospora, Lentzea, Nocardia, Nocardiopsis, and Amycolatopsis), and was repurposed for this study.

**Figure 2.**
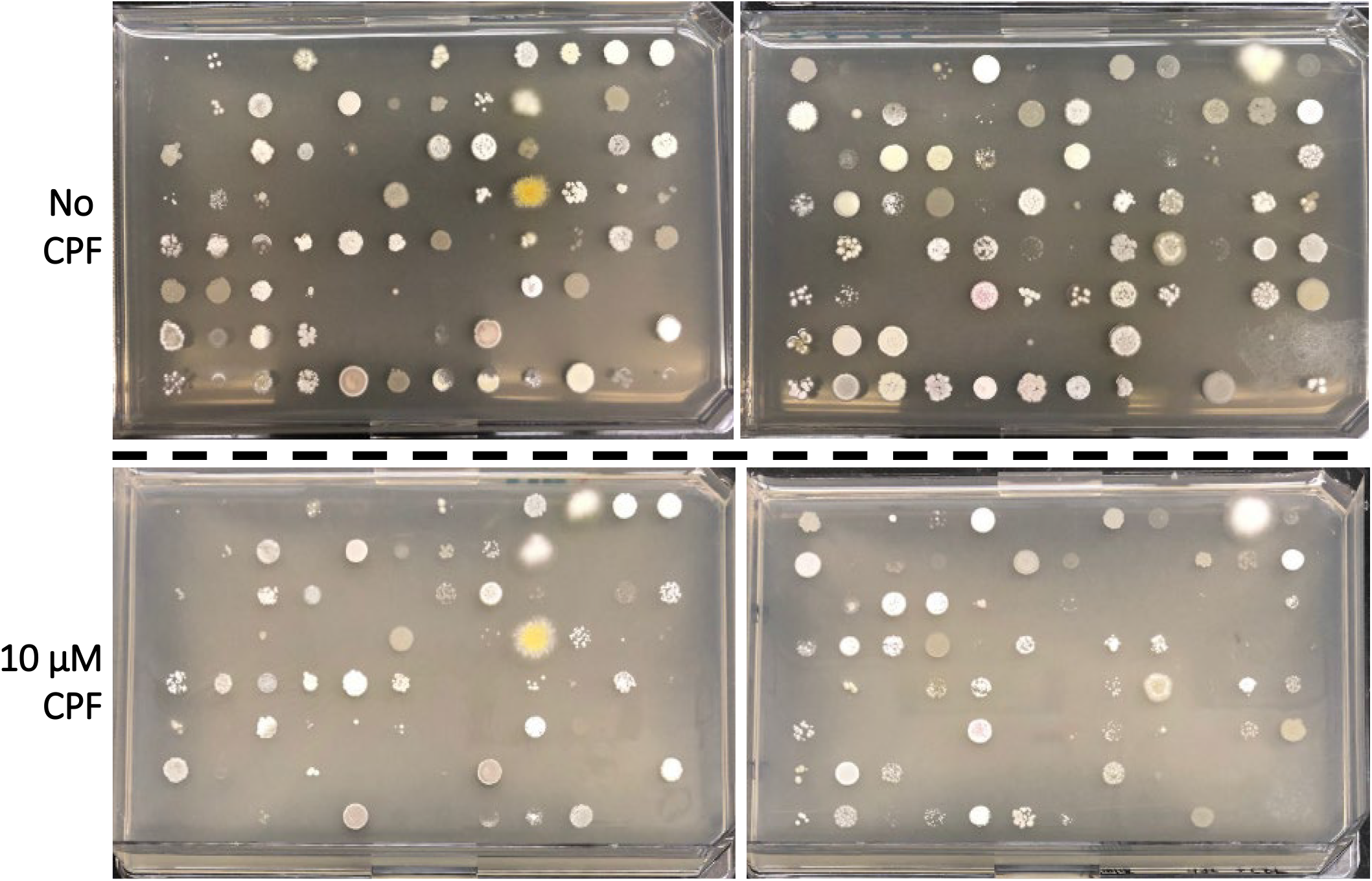
Screening the effects of chlorpyrifos exposure against a library of actinomycetes. A library of 196 actinomycetes were replica plated on media containing DMSO (Top) or 10 µM CPF (Bottom). Of the 196 strains plated 144 representative grew robustly on the DMSO control plates which served as the reference. The strains grown on CPF were compared to the counterpart on DMSO media.

### Chlorpyrifos exposure results in delayed growth of specific soil microbes

To eliminate the possibility that the inhibition we observed might be a secondary effect resulting from CPF-induced biosynthesis of cryptic secondary metabolites, a potential result of stimulation by exogenous chemical triggers (68, 69) or co-culture with other bacterial species (70), we next examined the effects of CPF on pure cultures of a panel of strains that were inhibited by CPF in the arrays. Utilizing select strains from the collection we determined the inhibitory effect of CPF by measuring colony diameter after a defined growth period. Spore preparations from four strains in the collection were plated on ISP2 agar plates containing either DMSO or 10μM CPF and incubated for 3 days at 28°C. Following incubation, colony diameters were measured and all tested strains exhibited a significant reduction in colony diameter upon exposure to CPF (**Fig. 3**). Specifically, strains B-2682, ECBC9.7, and QLW15 exhibited a 59.21%, 51.18%, and 72.6% decrease in colony diameter respectively, compared to these strains exposed to DMSO alone. Additionally, after the 3-d incubation period, there were no visible colonies from strain USNA 39.6B when plated on the CPF containing media (**Fig. 3**). We sequenced the 16S rRNA gene regions of the strains affected by CPF and have tentatively identified them as *Streptomyces griseus* B-2682, *Streptomyces paulus* ECBC 9.7 and *Streptomyces fulvissimus* QLW15.

**Figure 3.**
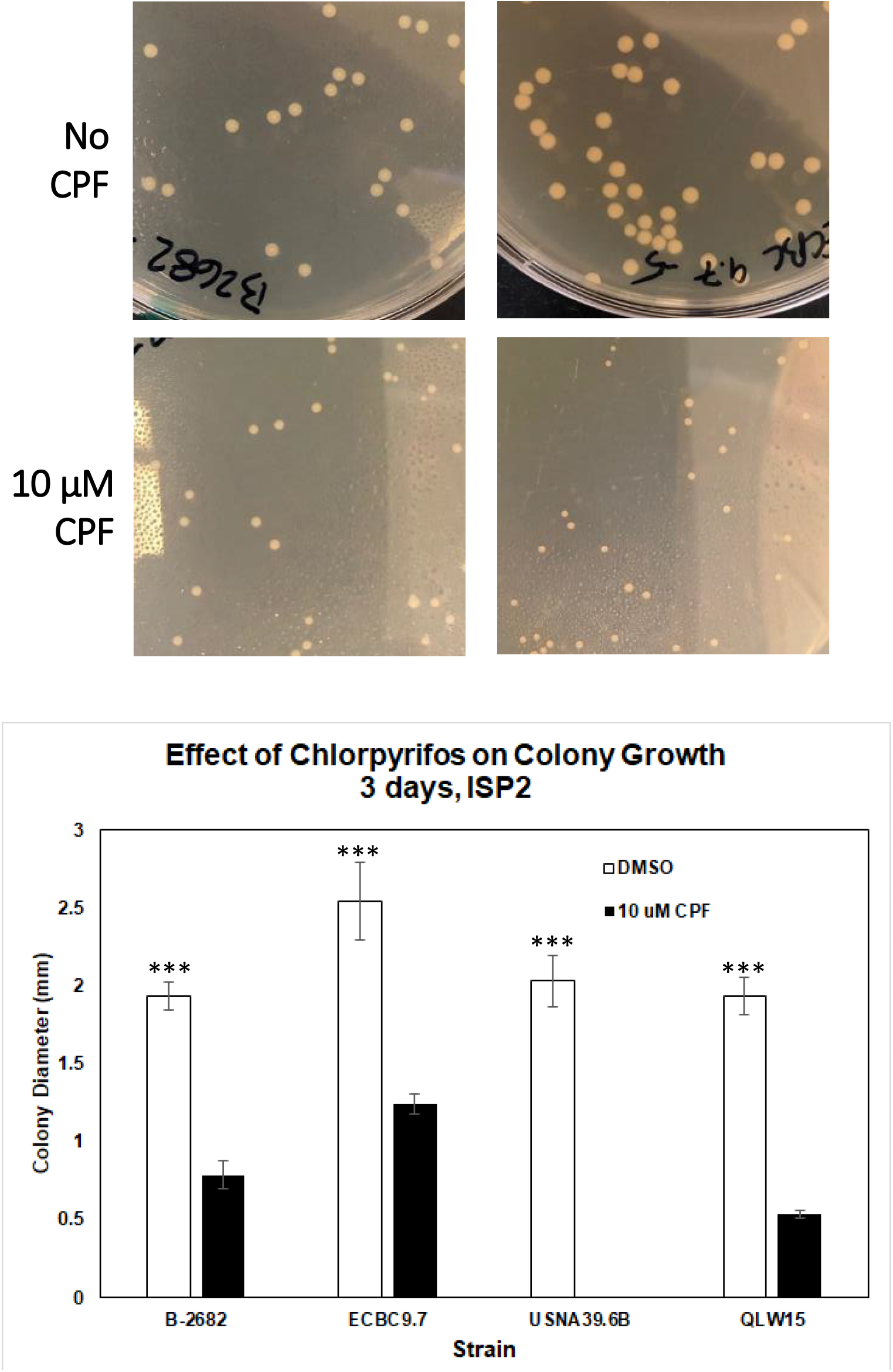
Effect of chlorpyrifos on colony size. Four Streptomyces strains were each inoculated onto an ISP2 agar plate with 10 µM CPF or DMSO and incubated for 3 days at 26°C. The diameter of a sampling of colonies was measured, represented (Top). The mean diameters for each strain tested were determined (bottom). Error bars represent SD. *** p<0.001 based on unpaired t-test between mean diameters from +/- CPF treatments.

### Actinomycete growth inhibition is specific to chlorpyrifos exposure

Having demonstrated that CPF inhibited a subset of organisms in our actinomycete collection, we next wanted to determine if these effects were observed with other OPPs. To test the effects of multiple OPPs, we conducted zone of inhibition experiments with the following compounds: (1) DMSO control, (2) CPF, (3) 3,5,6 trichloropyridinol, (4) malathion, (5) parathion, (6) monocrotophos, and (7) mevinphos (phosdrin). While not an organophosphorous compound, 3,5,6-trichloropyridinol (TCP) is one of the main degradation products of CPF upon hydrolysis, and previous reports have suggested that TCP has antimicrobial properties (49, 50, 71, 72). We speculated that the antimicrobial activities of TCP may explain the growth inhibition phenotypes we observed with CPF upon hydrolysis in the media. Whatman paper disks 6 mm in diameter were soaked with 10 μL of a 10mM stock solution for each compound and added to ISP2 plates with the indicated strain spread to form a lawn. The plates were monitored for growth and inhibition zones daily. We utilized *Mycobacterium smegmatis* as a well characterized actinobacterium control. We found that for each of the four strains tested, with the exception of *M. smegmatis* which was unaffected, CPF specifically inhibited growth whereas there was no observable effect from the alternative OPPs or TCP (**Fig. 4**). Additionally, similar to the observation on colony diameter, the effect of CPF was variable between strains tested, with some being more sensitive than others as indicated by the formation of larger zones of clearance or inhibition (**Fig. 4**). Across the strains tested here, at the given concentration, we did not observe any antimicrobial effects from TCP, suggesting that the inhibition associated with CPF is specific and not a result of degradation to TCP (**Fig. 4**).

**Figure 4.**
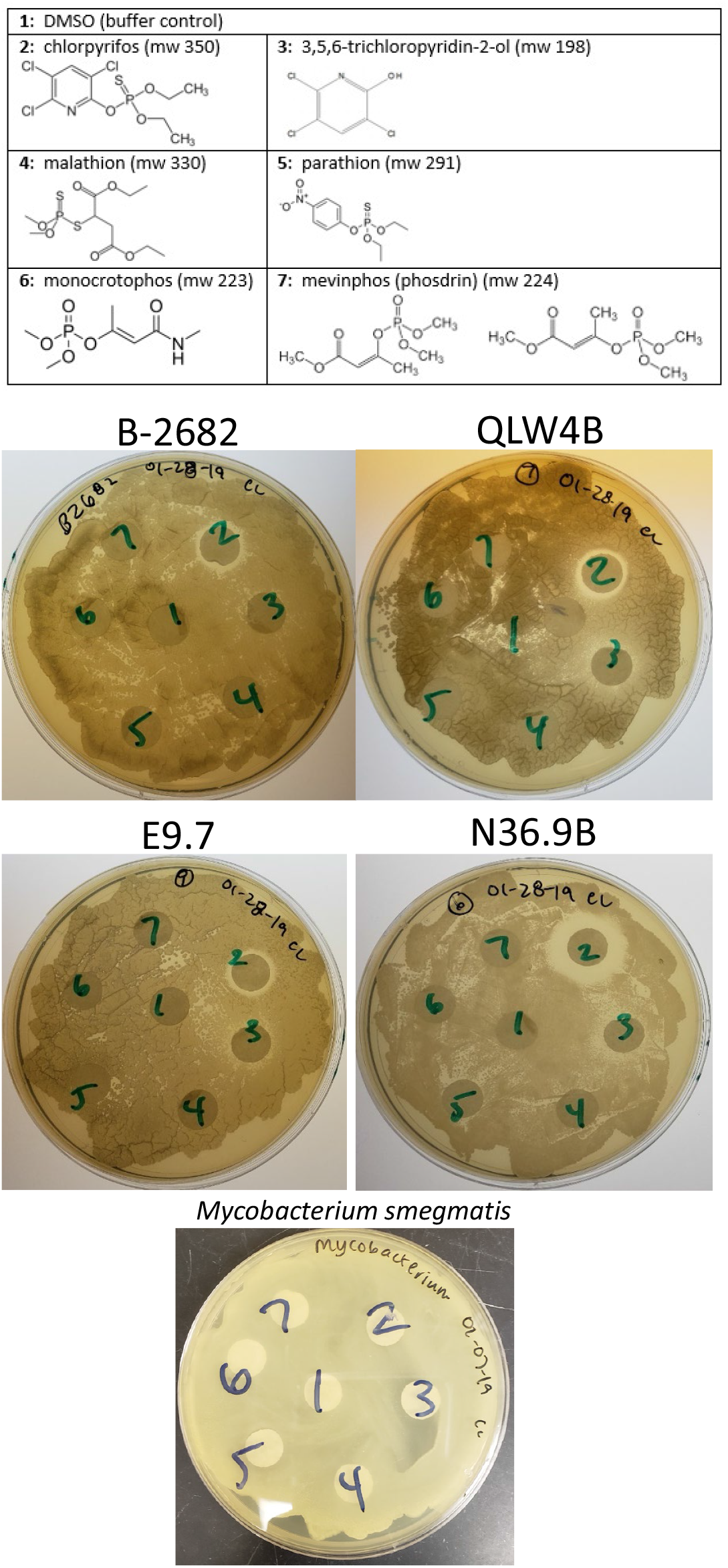
Determining the inhibitory effects of organophosphate pesticides on actinomycetes. Four Streptomyces strains and *Mycobacterium smegmatis* were each inoculated onto an ISP2 agar plate with discs dosed with 10 µL of 10 µM of (1) DMSO; (2) CPF; (3) TCP; (4) Malathion; (5) Parathion; (6) Monocrotophos; (7) Mevinphos and incubated allowing lawns to form. Following incubation zones of inhibition (ZOIs) were measured for each of the compounds tested.

We next wanted to determine whether the delayed growth in the presence of CPF was a phenotype observable across diverse species of bacteria. We conducted disk-diffusion assays with the same panel of OPPs tested against the actinomycetes with representative bacteria: *Escherichia coli, Bacillus subtilis, Bacillus thuringiensis, Staphylococcus epidermidis* and *Burkholderia thailandensis*. Following incubation, none of the compounds at the given concentration exhibited any deleterious effects to growth against the strains tested here (**Fig. 5**). Taken together these data demonstrate that, at the tested concentrations of OPPs, only select actinomycetes are specifically susceptible to CPF exposure.

**Figure 5.**
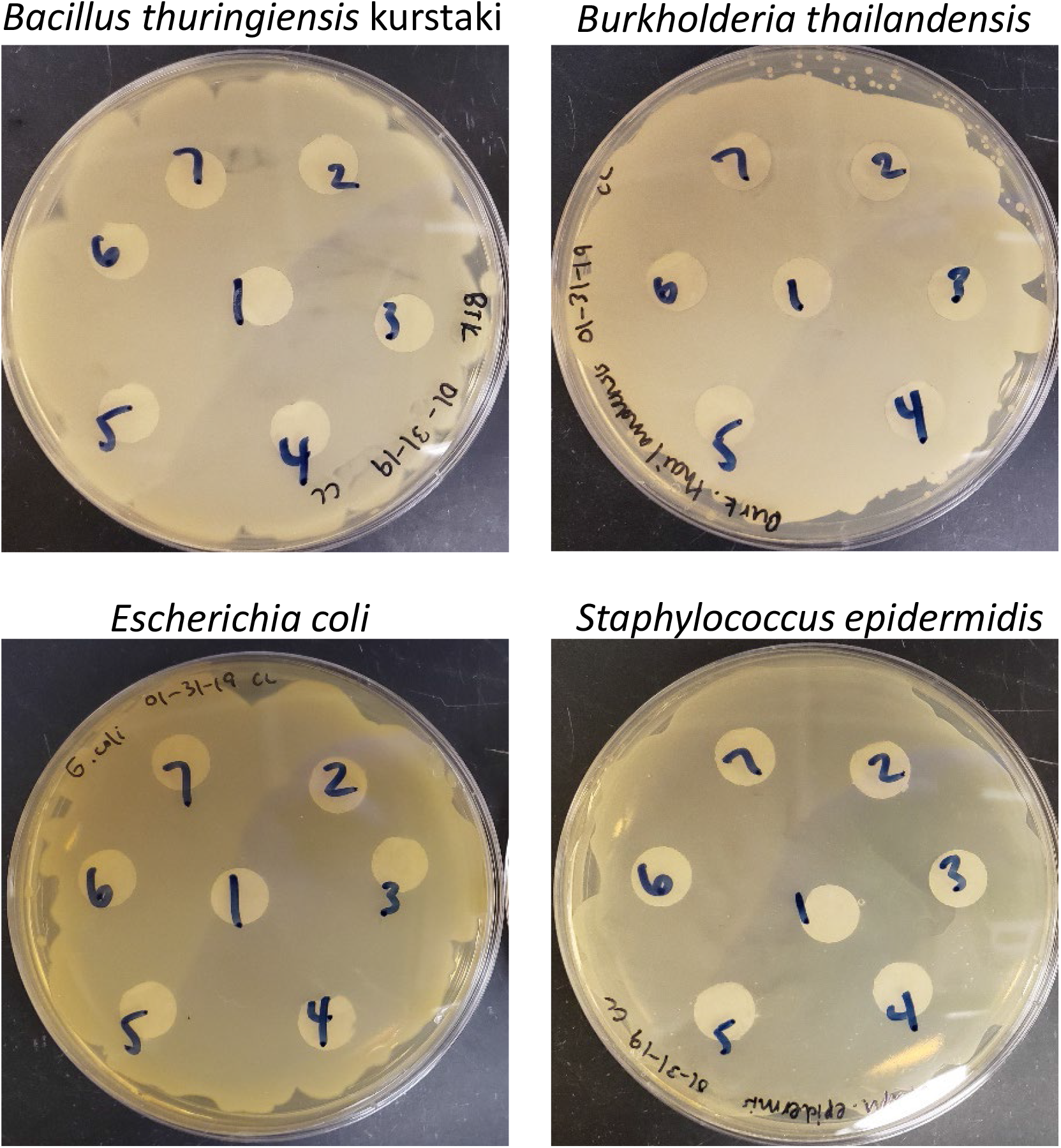
Determining the inhibitory effects of organophosphate pesticides on non-actinomycetes. Four representative bacteria: *Bacillus thuringiensis* serovar kurstaki, *Burkholderia thailandensis, Escherichia coli* and *Staphylococcus epidermidis*, and were each inoculated on appropriate media with discs dosed with 10 µL of 10 µM of (1) DMSO; (2) CPF; (3) TCP; (4) Malathion; (5) Parathion; (6) Monocrotophos; (7) Mevinphos and incubated allowing lawns to form. Following incubation zones of inhibition (ZOIs) were measured for each of the compounds tested.

### Species and strain specific differences in chlorpyrifos sensitivity

Since many strains in our own collection have not been well characterized, and all of the identified strains that showed CPF-sensitivity belonged to the genus *Streptomyces*, we sought to better understand the taxonomic breadth of inhibition by testing a selection of soil associated actinomyces species/strains obtained from the Agricultural Research Service Culture Collection (NRRL) for CPF sensitivity (**Table 1**). The collection was chosen to represent phylogenetically diverse actinomycetes found in various soil environments. The collection was evaluated for CPF-induced sensitivity via the zone of inhibition test comparing 10 μM CPF or DMSO control. Similar to the observations described above, we found that CPF had variable inhibitory effects towards some isolates in our NRRL panel (**Table 1**). In general, the phenotypes could be classified as either a large zone of inhibition (ZOI), a small ZOI, growth up to but not on the CPF disk, or completely unaffected by CPF (**Table 1**). In total, 8 of the 19 strains tested (42%) exhibited measurable ZOI upon CPF exposure (**Table 1**). The most dramatic example of CPF sensitivity was observed in *Micromonospora chokoriensis* for which we measured a ZOI of 40.5 mm (**Table 1**). This particular strain of *Micromonospora chokoriensis* was isolated from soy bean roots in Illinois, USA. This isolation source is particularly relevant as soybeans are one of the main target crops for CPF application (16). In screening our panel of representative actinomycetes, there are differences in CPF sensitivities between highly related bacterium, such as *Kibdelosporangium aridum* subspecies *aridum* which exhibited a ZOI of 12.2 mm and *Kibdelosporangium aridum* subspecies *largum* which was unaffected by CPF exposure (**Table 1**). Collectively, these results align with those from our internal collection in that the inhibition of growth upon CPF exposure only effects select actinomyces and differences in sensitivity exist between related strains/species.

**Table 1:**
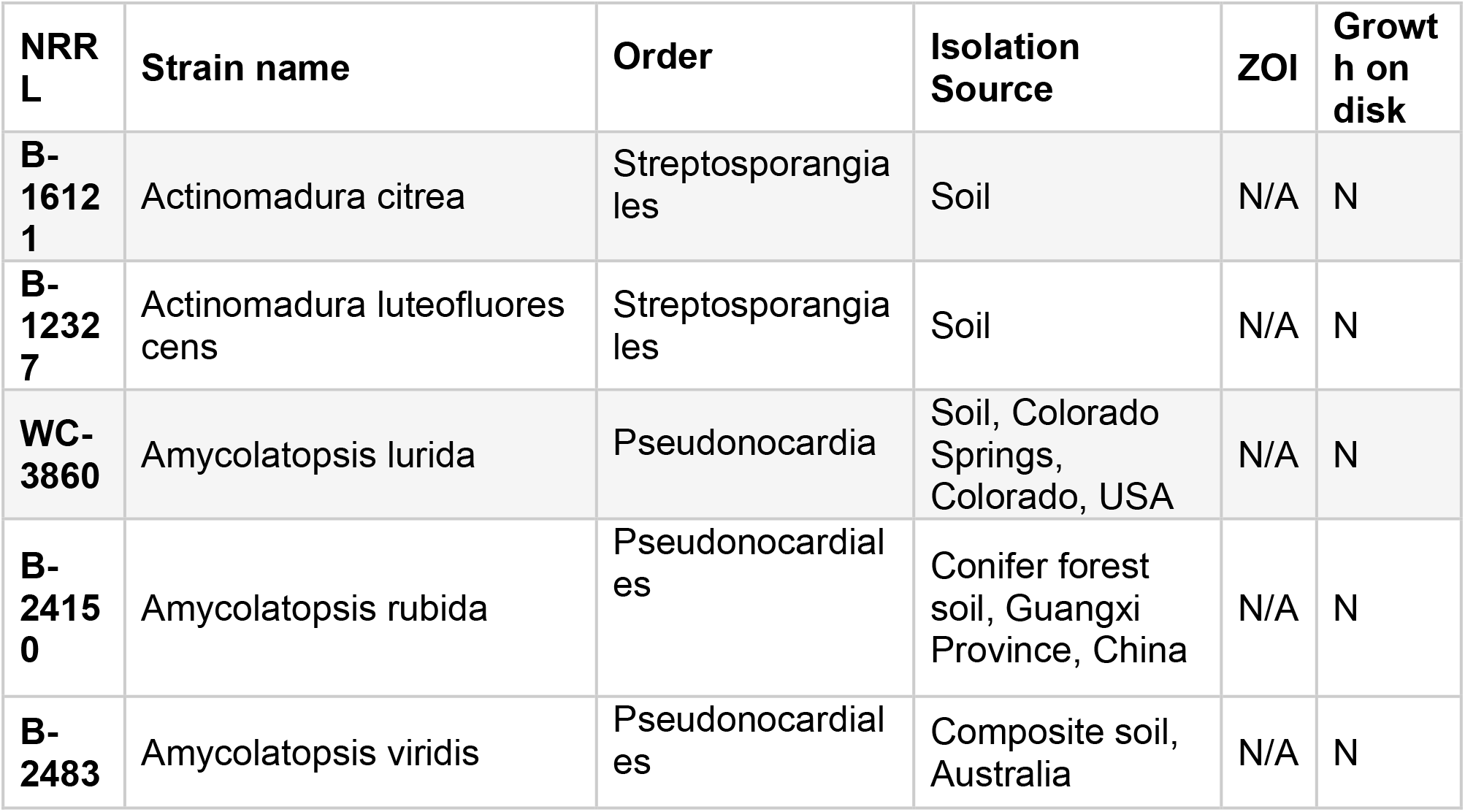
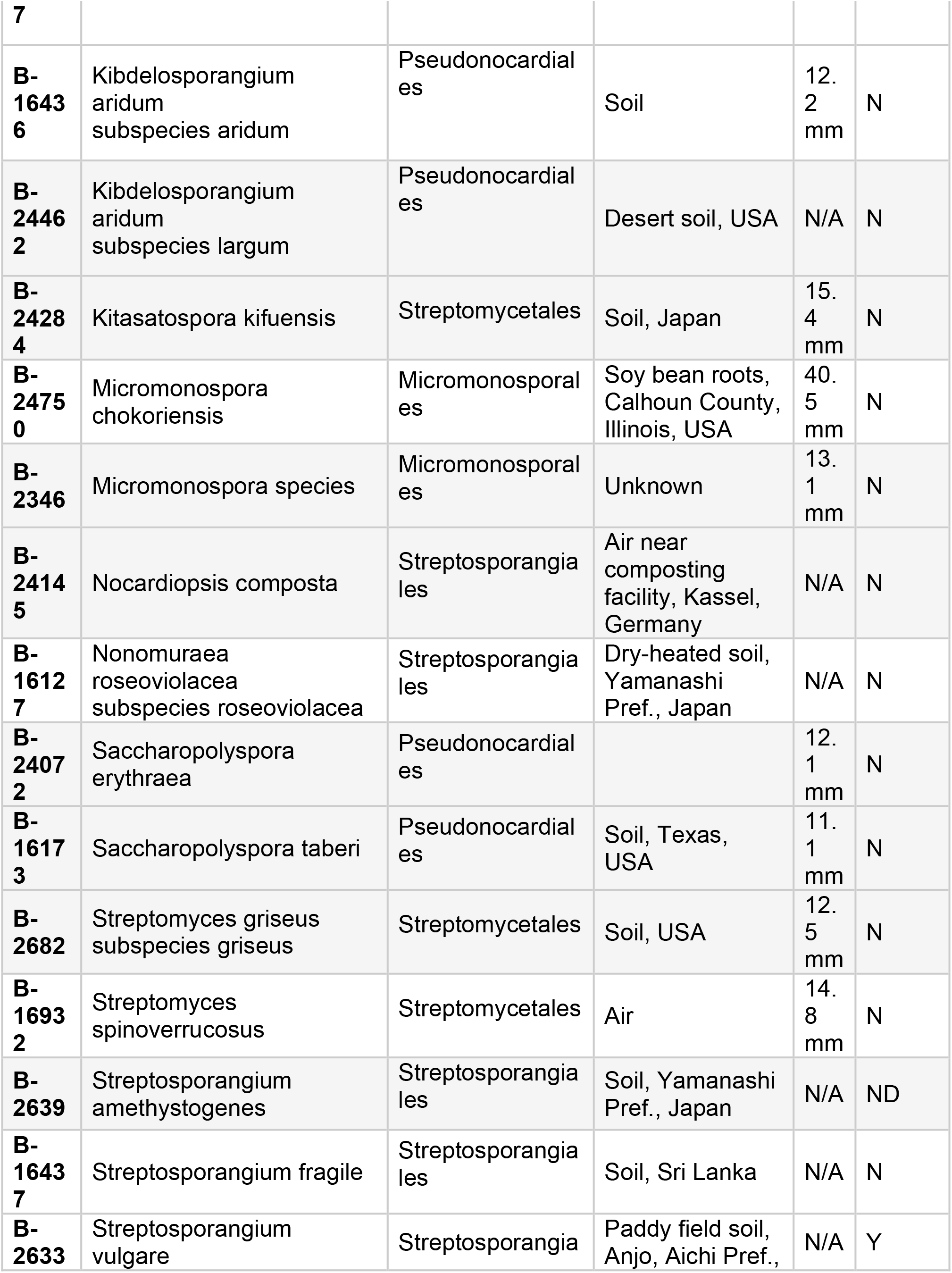
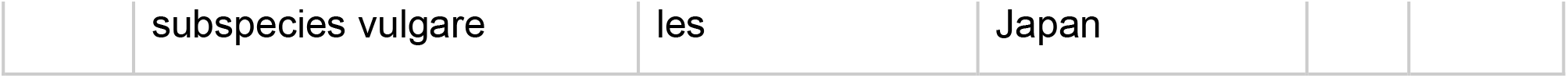
Evaluation of NRRL representative actinomycetes exposed to CPF

## Discussion

With this work, we set out to characterize the impact of OPPs on soil microbes. This class of insecticides has been highly scrutinized because of its deleterious effects on non-target organisms such wildlife and humans, particularly agricultural workers. Despite these concerns associated with OPPs, millions of tons are used annually in agriculture systems worldwide, ultimately making their way into the food supply (73). While the risks to humans are documented, much less is understood regarding the potential for OPPs to interact with the soil microbiota. As an essential component of agricultural ecosystems, alterations in soil microbe populations may result in deleterious downstream outcomes. Therefore, we set out to specifically determine how OPPs affect the growth of soil actinomycetes.

We found that when specific actinomycetes are exposed to chlorpyrifos at agriculturally relevant concentrations, there is significant growth inhibition and, in some cases, strains were completely unable to grow. While the *in vitro* concentration we selected (10 μM) assumed full solubilization of the compound, the solubility of CPF in water is 1.4 mg L^-1^ or 4.0 μM (74). Therefore the actual exposure to CPF *in situ* may vary, the compound can adsorb to soil particles due to its high soil partition coefficient (75). Given that streptomyces species in particular display hydrophobic structures that protrude outside of aqueous environments and potentially in direct proximity to soil particles (65, 76-80), it is possible that the mode of interaction with the pesticide in these experiments may not represent *in situ* exposure routes. We speculate that the hydrophobic nature of the aerial hyphae, which are expressed within and between soil particles (81, 82) may allow relatively insoluble CPF and other pesticides to partition to the surface structures of actinomycete species, and potentially increase access to cell-surface targets.

The mechanism(s) by which CPF inhibits the growth of sensitive strains is unknown. OPPs and similar compounds can inhibit a variety of non-target enzymes, particularly serine proteases and esterases. In separate activity-based protein profiling studies, analogs of CPF and the nerve agent VX were shown to bind and form adducts with several molecular targets in addition to cholinesterases (83-85), including enzymes involved in the TCA cycle. Furthermore, the organophosphate reagent diisopropylfluorophosphonate is commonly used as a non-specific serine protease inhibitor. The isolation of resistant or hypersensitive mutants and identification of bacterial CPF-binding enzymes should illuminate the mechanism by which CPF blocks growth or the mechanisms by which isolates resist inhibitory action of CPF.

It is important to note that the phenotypes observed in this study also do not take into account potential effects of CPF on the complex microbial communities in which these strains exist in soils. The complexity of the soil microbiome may offer protection from the toxic effects of OPPs through mechanisms such as biodegradation or by providing protected niches within single- or multispecies biofilms. Previous investigations examining the impact of pesticides on microbes have primarily focused on dynamic population shifts and overall enzyme activity, and in fact there have been several conflicting reports regarding how microbial populations are affected(51, 56, 63, 86-88).

In our experiments, the growth inhibition was specific to certain strains of bacteria while others remained unaffected by CPF. A recent study analyzed the effects of CPF and malathion on microbes and enzyme activity in soil collected from Punjab, India. This study found that in soil containing CPF and malathion at concentrations ranging from 10-250 ppm there was no significant loss in total microbial populations or actinomycetes specifically (63). Comparatively, in our studies, at lower CPF concentrations (3.5 ppm), we observed species/strain specific inhibition of actinomycetes. These findings would suggest that at coarser levels of taxonomic resolution, there may not be observable changes in population, while changes at the species level – such as selective inhibition by CPF – would be missed.

The data presented have demonstrated that there are members of the actinomycete family which are CPF resistant and others which are inhibited by CPF. The CPF induced inhibition of select microbes decreases competition for the resistant population, allowing for niche expansion. Supporting the potential for possible niche expansion following CPF exposure, there are several examples of actinomycete strains that are capable of degrading CPF and utilizing the pesticide as a sole carbon source (47, 57). In the environment, it is possible that CPF is inhibiting select actinomycetes while others are resistant or even growing on CPF thereby shifting community-level dynamics within the group.

Our studies indicated that the inhibition observed was due specifically to CPF as we did not observe any effects from chlorpyrifos’ main degradation product or several other OPPs tested. Previous studies have demonstrated that malathion at 50 ppm decreased actinomycete populations in soil by 37% in one week following application (89). The absence of growth inhibition with malathion observed in this study could be related to the low concentration used here (only 3.5 ppm). It is possible that higher concentrations of malathion could lead to growth inhibition.

In the environment, CPF readily hydrolyzes to the primary degradation product 3,5,6-trichloropyridinol (TCP), which has its own associated risks and health concerns (37, 90, 91). Numerous biodegradation studies, focused on utilizing microbes to remediate CPF found that the accumulation of TCP hindered microbial growth (49, 50, 71, 72). We speculated that CPF-mediated inhibition of actinomycetes was due to hydrolysis to TCP in the media. We found that, at the concentrations tested, TCP did not inhibit any of the strains which were sensitive to CPF demonstrating that the inhibition was specific to CPF. A previous study demonstrated that the chlorpyrifos oxon, an intermediate degradation metabolite, was ∼26 times more inhibitory than CPF towards soil microbes (88). It is possible that the growth phenotypes observed in this study are a result of CPF oxon interactions, which should be explored in further detail.

There is likely no “magic bullet” pesticide which exhibits no off-target effects while providing the benefits needed to support agricultural needs worldwide. This study sheds light on how the prevalent organophosphate pesticide CPF can selectively exhibit antimicrobial effects on actinomycete subpopulations. Finding that some actinomycetes are unaffected by CPF suggests that there are genetic elements and/or associated phenotypes present within these strains that confer resistance to CPF. One possible explanation for CPF resistance is the presence of hydrolases which can degrade CPF in the environment prior to the molecule accessing the cytoplasm. A second possibility is that the low solubility of CPF prevents it from crossing the inner membrane in resistant strains whereas sensitive strains may specifically or nonspecifically transport CPF into the cell. It is not clear why CPF is the active compound against the sensitive strains here and not TCP which has previously been reported to exhibit antimicrobial activity. Previous studies have shown that at sub acetylcholinesterase inhibitory concentrations, CPF can inhibit non-target hydrolases in mammalian tissues (85, 92, 93). It is possible that CPF is inhibiting essential hydrolases in the sensitive actinomycetes resulting in growth defects. Further investigations should be directed at illuminating the mechanism of action for CPF growth inhibition in these organisms. While total actinomycete population counts may not necessarily be changed upon CPF exposure, this study highlights that CPF exposures most likely alters the taxonomic composition of soil actinomycete communities.

## Materials and Methods

### Collection of the actinobacterial strain library

The actinomycete library was assembled from organisms isolated from air samples. The details of actinobacteria isolation and detailed analysis of the collection composition will be published elsewhere (R. Killens-Cade *et al*., in preparation). Briefly, strains were collected and isolated onto Humic Acid Vitamin Agar or Kings B Medium using a SASS-180 single-stage impact sampler (Bioscience International, Rockville, MD, USA) or an MAS-EC0100 (EMD Millipore, St. Louis MO) using published methods (94). Colonies exhibiting characteristic actinomycete morphology (e.g. substrate mycelia, aerial hyphae, and/or sporulation) were streaked to homogeneity, sporulated on solid media, and stocked using standard methods (95).

### Strains and culture conditions

The actinobacteria were routinely cultured from frozen spore preparations via inoculation into Difco International Streptomyces Project (ISP) medium 2 (ISP2) (Becton, Dickinson and Company, Spark, MD) and incubated at 28° C. The actinobacteria listed in table 1 were obtained from ARS NRRL culture collection (https://nrrl.ncaur.usda.gov/cgi-bin/usda/index.html) and were selected to include a broad representation across the actinobacteria. When pesticides were included in the growth medium, they were diluted from a 10 mM stock solution in DMSO directly to the growth medium after sterilization.

### Replica plating soil microbe library

Frozen spore preparations were used to inoculate 1 ml volumes of International Streptomyces Project-2 (ISP2) medium in deep 96-well plates (Enzyscreen BV, Heemstede, The Netherlands) and incubated at 28°C at 300rpm for 96 h after which most cultures became turbid (67). The outgrowth was then used to replicate the libraries on International Streptomyces Project-4 (ISP4) agar containing either 10μM CPF or DMSO control and incubated for 3 days at 28°C.

### Disk-diffusion studies using organophosphate pesticides

Single colony picks from ISP2 plates were added to 1 mL 0.1% PBS-Tween80 for spore-forming organisms. Non-actinobacteria bacteria inoculations were prepared with a single colony into 5 mL of LB broth and shaken at 37°C for about 5 hours, until turbid. Actinobacteria were spread onto ISP2 agar, non-actinobacteria were spread onto Mueller Hinton agar, at 400 μL for each plate. Once absorbed, 6-mm diameter Whatman paper disks soaked in 10 μL of 10 mM pesticide in dimethyl sulfoxide (DMSO) or in a solvent-only control were placed on the agar surface and transferred to the incubator for growth. Actinobacteria were grown at 28°C, non-actinobacteria at 37°C, and monitored for growth daily. Plates were removed from the incubator once lawns were observed.

## Acknowledgements

Funding was provided by the Defense Threat Reduction Agency. This research was performed while N.D.M held an NRC Research Associateship award at CCDC-CBC. We thank Soumia Bekka and Andrew Marinich for assistance with environmental sampling at CCDC-CBC.

